# Ab-AMR: A Comprehensive Repository of *Acinetobacter baumannii* to Understand the Molecular Landscape of Antimicrobial Resistance

**DOI:** 10.1101/2022.07.17.500328

**Authors:** Tina Sharma, Rakesh Kumar, Anshu Bhardwaj

## Abstract

Ab-AMR is a comprehensive repository of drug resistance mechanisms in *Acinetobacter baumannii*. The current version of Ab-AMR provides a drug resistance profile of 788 genomes. In order to ensure that the datasets in Ab-AMR have relevance both to the research and clinical community, standards of defining MIC breakpoints, whole genome sequence quality metrics as defined by EUCAST/CLSI and classification of isolates into susceptible, MDR, XDR and PDR as defined by CDC/ECDC are implemented. As of now, 364 DR determinants associated with antibiotic inactivation (β-lactamases, aminoglycoside modification, chloramphenicol acetyltransferase), efflux, protein modulating permeability and alteration of target site are comprehensively annotated. In addition, data from pangenome analysis across 788 genomes is also provided for identification of core and accessory DR determinants. *AB* ATCC 17978 (Accession-CP000521.1), the reference strain, was annotated on January 1, 2014 but subsequently the same strain was re-annotated on March 21, 2017 (NZ_CP018664.1) due to incorrect assembly. Therefore, the genome comparison of both, 2014 and 2017 versions were performed for maintaining the correct annotations as most of the literature data referred to the earlier version of the reference genome. In Ab-AMR, the reference coordinates of the revised reference genome are used to represent manually curated and comprehensively annotated data on 614 essential genes, 1334 genes mapped to pathways, 221 PDB structures, 81 reported drug targets, 364 genes with reported resistance mechanism, 118 transcription factors, 4 sigma factors and 14 two component systems. Ab-AMR is made using the standard php-mysql framework and offers various search tools including a query builder that facilitates query on over 60 different features for addressing complex questions like core genes which are also essential and have a role to play in drug resistance with no known human homolog, etc. Ab-AMR offers a centralized data resource for systematic mapping of DR determinants, both plasmid and chromosomal mediated, along with deep annotation of clinical isolates.

Database URL: https://datascience.imtech.res.in/anshu/ab-amr/

## Introduction

In recent years there is an unprecedented increase in high throughput DNA sequencing of genomes and metagenomes towards bacterial typing, molecular epidemiology investigations, and in-depth pathogenetic studies. In addition to identification of new markers to detect microbial infections, whole genome sequencing (WGS) can provide insight into the role of genomic variation in strain evolution and its impact on transmission in real time. In addition, it is also shown that the drug-target interaction is far more complex than previously anticipated with a single antimicrobial acting via interactions with several proteins in the pathogenic organisms. This also translates to a wide array of mechanisms that can lead to drug resistance. Resistance to several antibiotic classes lead to multidrug resistance (MDR), extensively drug resistance (XDR) and pan-drug resistance (PDR) strains (1–4). Understanding the dynamics of MDR, XDR and PDR pathogens requires consolidating drug resistant determinants of various clinically resistant phenotypes along with detailed understanding of their role in drug resistance (5). In case of *A. baummanii* it has been observed that it has an open pan-genome (performed on 349 strains) (6), which clearly indicates that sequencing of more strains of AB can lead to identification of novel drug resistance (DR) determinants. As has been shown recently, genome sequencing of a novel strain of AB viz. DMS06669 led to identification of eight new DR determinants (7). In this study, two antibiotic resistant genes (blaNDM-1 and blaOXA-58) were reported in a single strain for the first time. Similarly, several other studies have been performed over years that led to identification of novel drug resistant determinants and mechanisms of resistance (8). There has been a consistent rise in reporting of DR determinants in AB as evident from keyword-based search in PubMed. However, this information is scattered across literature and it is not feasible to comprehend the complexity of emergence of resistance in AB for several different antibiotics. Therefore, it is imperative that a consolidated platform is created not only to provide a comprehensive assessment of known drug resistant determinants but also to guide discovery of new ones.

A number of resources for the identification of resistome have been developed over years. For instance, Antibiotic Resistance Genes Database (ARDB) (9), which provides information on antibiotic resistance and facilitate annotation of resistance determinants in newly sequenced organisms. ARDB was last updated in July 3, 2009 (https://ardb.cbcb.umd.edu/) and is now integrated with Comprehensive Antibiotic Resistance Database (CARD) (10,11). As of April 06, 2021 CARD contains resistome predictions for over 200 pathogens (https://card.mcmaster.ca/resistomes). It has 491 protein homolog models, 3 protein overexpression model, 4 protein variant model and 1 rRNA gene variant model for AB for 31 classes of antibiotics (accessed on April 06, 2021-CARD prevalence data from https://card.mcmaster.ca/) (10). Similarly, ResFinder 4.0 (12) provides information on acquired resistance genes for various pathogens except AB. The Pathosystems Resource Integrated System (PATRIC) (13) provides resistance gene data from ARDB, CARD and National Database of Antibiotic Resistant Organism (NDARO) (14), PATRIC reports 24 drug resistant genes in AB based on CARD (21 genes) and ARDB (3 genes) databases. Despite these centralized efforts, several reported drug resistance determinants are not available in these resources. For example, Trm, a *trm* disruption which is associated with tigecycline-resistant strains of AB (15); Abrp, belongs to peptidase C13 family and is shown to be responsible for decreased susceptibility towards several antibiotics (tetracycline, minocycline, doxycycline, tigecycline, chloramphenicol, and fosfomycin) (16); CraA, which is a major facilitator superfamily efflux pump, is shown to confer chloramphenicol resistance (17); several cell division proteins like zipA, zapA, and ftsK (18) confer increased β-lactam susceptibility etc. are not present in PATRIC and CARD databases.

So far, species-specific drug resistance databases have been developed only for *M. tuberculosis* (Tuberculosis Drug Resistance Mutation Database (19), MUBII-TB-DB (20)) and for *E.coli* u-CARE (21). For understanding drug resistance mechanisms in *Mtb* and *E.coli*, these databases were considered to be indispensable (22). Therefore, in this study an AB centric repository, Ab-AMR, is developed.

The strain ATCC 17978 is a clinical isolate from 1951 originally named *Moraxella glucidolytica nonliquefaciens*. Its genome was sequenced in 2007, is 3,976,746 bp with 3830 ORFs. The genome contains multiple pathogenicity islands that encode genes associated with virulence, such as Type IV secretion systems, drug and heavy metal resistance proteins, fimbriae, and mobile elements (23). *A. baumannii ATCC 17978* has been considered to be the best and first studied strains of Acinetobacter species, and is a useful strain to study the pathogenesis of other strains due to its intrinsic sensitivity to almost all antibiotics used to treat AB infections (24,25). In 1951, *A. baumannii ATCC 17978*, was obtained from a case of fatal meningitis for which the antibiotic resistance profile was not previously reported (26). The discovery of *A. baumannii ATCC 17978*, in early 1950s i.e., before the discovery of latest β-lactamase, cephalosporins, glycopeptides and macrolides make this genome sequence a representative genome for comparative genomics of AB (27). In the 1970s, this organism was susceptible to most antimicrobial agents except sulphonamides (28). AB has now become a major cause of hospital acquired infections all over the world due to its remarkable ability to acquire resistance determinants to various kinds of antimicrobial agents (29). Since 1970s, the emergence of multidrug-resistant (MDR) *Acinetobacter baumannii* strains among sick and immune-compromised patients have become a serious concern for health-care (30). *Acinetobacter baumannii ATCC 17978* (Accession-CP000521.1) was annotated on January 1, 2014 and subsequently latest re-annotated on 2015 (NZ_CP012004.1, latest genome modification date is February 8, 2021) and March 21, 2017 (Accession-NZ_CP018664), due to the presence of a 148 kb conjugative plasmid, pAB3, fragments of which erroneously merged into the chromosome in the original 454-based assembly (Accession ID: CP000521.1) (31). The genome sequence of ATCC 17978 i.e., Accession-CP000521.1 with 3976746 bp was released on March 1, 2007 and sequence at NCBI was last modified on January 31, 2014. Most of the annotations among literature refer to this original version (Accession-CP000521.1) of ATCC 17978 which is still used as a baseline for comparative genome analysis of AB. A paper published in “September 26, 2019 (31) clearly mentioned that the original version i.e. CP000521.1 was sequenced using 454 pyrosequencing technology and assembled using PCR. Another version of ATCC 17978 i.e., CP012004.1 was available at NCBI named *Acinetobacter baumannii strain ATCC 17978-mff* chromosome released on July 14, 2015. The sequencing technology used to sequence this version was Illumina, Pacbio and which was available with 3857743 bp. The sequencing of this genome revealed the presence of a 148 kb conjugative plasmid, pAB3. The original genome sequence for *A. baumannii ATCC 17978* i.e., Accession-CP000521.1 had two plasmids (pAB1 and pAB2). However, it was subsequently revealed that ATCC 17978 carries a third plasmid (pAB3) that was incorrectly assembled as chromosome sequence in the first assembly. Later, on December 12, 2016, NCBI released different version of the ATCC 17978 i.e., NZ_CP018664 with genome size 4004792 bp and the sequencing technology used was Illumina MiSeq and the source of isolation was blood. The ATCC genome portal which is known to have a very high quality of reference strain has ATCC 17978 reference strain with 4 contigs comprising 4,075,779 bp https://genomes.atcc.org/genomes/e1d18ea4273549a0. In our analysis, we used the annotation of the latest version i.e., NZ_CP018664 (2016) and we ensure that the gene annotation is significantly aligned with the chromosomal genome, not the plasmid.

Ab-AMR is designed to provide a comprehensive assessment of drug resistant determinants for ~800 AB strains for which whole genome sequence data is available. In order to ensure that the data across clinical isolates is comparable, three international standards were used, namely, CLSI for MIC breakpoints, CDC/ECDC for phenotype classification and EUCAST for assessing genome quality of sequenced clinical isolates. In Ab-AMR, comprehensive annotation of the whole genome sequences is performed to map the known drug resistant determinants in AB from several resources including manual curation from literature. These DR determinants are then evaluated with respect to the pan genome of AB as well as their presence on chromosomes or plasmids and functional assignment to mechanisms of drug resistance. Other supporting annotations such as availability of crystal structure, pathway mapping, essentiality, drug targets, etc. is also provided. Comprehensive annotation of AB makes Ab-AMR a central resource for in-depth understanding of drug resistance at molecular level for existing as well as newly sequenced genomes of AB. An integrative approach to prioritize potential drug targets was also implemented. The data in Ab-AMR is represented in a structured manner as a web interface for easy access and search and is designed based on FAIR principles.

## Material and Methods

### 1. Data source and data preparation

Genome sequences for Ab-AMR are mostly curated from PATRIC with few genomes included from literature. Several filtering criteria like human host, good quality genomes (including good genome quality metrics by EUCAST), MIC standards (CLSI/EUCAST), etc including removal of incomplete entries is applied in PATRIC. From over 4000 genome sequences available, 788 genomes are shortlisted for further analysis.

#### 1.1 Deep annotation of the reference genome (NZ_CPO18664)

All the 788 genomes were annotated using RASTtk release 1.3 (https://github.com/TheSEED/RASTtk-Distribution/releases) with an options as - scientific name (*Acinetobacter baumannii*), domain (bacteria) and genetic code (32).

#### 1.2 Genome comparison of Accession-NZ_CP018664

The mapping of the reference strain of AB (Accession - CP000521.1) was performed with the revised reference genome AB *ATCC 17978* (Accession NZ_CP018664) due to the presence of a 148 kb conjugative plasmid, PAB3, fragments of which erroneously merged into the chromosome in the original 454-based assembly. Therefore, Megablast was used to map the coordinates of the genes from the newer version of the genome with the older version of the genome with 28-word size (default), >= 98% query coverage per subject, >=98% query coverage per HSP, >=98% percentage identity. Once the revised reference genome is mapped, it is comprehensively annotated with respect to gene function, pathways, DR determinants, pan-genome, etc using RAST, Blast2Go, KEGG, BioCyc, DEG, PATRIC, CARD, PRAP, BPGA, Roary, etc. Extensive data mining is also performed for fine annotation of the reference genome. Since, the literature contains the information of older versions of the genome, we mapped and revised the annotated data for ATCC 17978.

**Link**: *https://github.com/tinabioinfo/Comparision_of_reference_genome/blob/main/cover_ol_genes(2).pl*

### 2. Data mining from literature

To retrieve data on drug resistant determinants from literature, PubMed is searched using keywords like “((resistant gene) AND (*Acinetobacter baumannii*))” and the gene name or protein accession ID were retrieved from literature and the sequences were retrieved from NCBI or batch Entrez for further analysis. These sequences were then mapped to the selected reference genome (NZ_CP018664) and 787 clinical isolates selected for Ab-AMR. Literature was also thoroughly searched for manual curation of drug targets, transcription factors, sigma factors and two-component systems (2000-2018). pubmed.mineR(33) (https://mran.microsoft.com/snapshot/2017-02-04/web/packages/pubmed.mineR/index.html) was used to retrieve research papers using different keywords for instance ((*Acinetobacter baumannii*) AND targets) AND drug targets-2000-2018 (till September 4, 2018).

### 3. Retrieving annotations from databases

Several databases were referred to systematically annotate the AB reference genome. DR determinants of AB were retrieved from CARD’s prevalence data and protein homolog models and PATRIC (https://www.patricbrc.org/) in addition to literature. CARD prevalence data for DR determinants, is generated using the Resistance gene identifier (RGI) tool. DR determinants from PATRIC were downloaded from the specialty genes tab and applied filters for “Antibiotic resistance” and “Literature”. For annotation of essential genes in AB, DEG (http://www.essentialgene.org/) was referred. Drug targets reported in AB were searched in PATRIC and DrugBank (https://go.drugbank.com/). The pathways for reference genome were retrieved from Biocyc (https://biocyc.org/), KEGG (https://www.genome.jp/kegg/) and PATRIC (https://www.patricbrc.org/). RCSB PDB (https://www.rcsb.org/) and Uniprot (https://www.uniprot.org/) were referred for PDB structures. It was observed that not all DR determinants reported in literature were mapping to the reference genome. Therefore, to obtain pathway annotation for these DR determinants, BLASTKoala was used to get KO (Kegg ontology) numbers by uploading the amino acid sequences of DR determinants, with taxonomy group “470” and species_prokaryotes in KEGG genes database. The DR determinants were further comprehensively annotated for known resistance mechanisms with mutations conferring resistance (from literature and CARD) and similar resistance mechanisms reported in other species along with locus tag, product, gene ontology etc.

### 4. Pan genome analysis (788 clinical isolates)

Pan-genome analysis of 788 clinical isolates was carried out using a bacterial pan-genome analysis pipeline (BPGA)(34) which uses Usearch (v10.0.240_win32) for the clustering of the orthologous genes with 50% identity. The estimated core, accessory (dispensable) and unique (strain specific) genes of AB were then aligned to DR determinants, drug targets and essential genes with 50% percentage identity cut off and >= 90% query coverage with 0.0001 e-value.

### 5. Ab-AMR use case: Drug target prioritization

Human proteome was downloaded from (‘ftp://ftp.ncbi.nlm.nih.gov/refseq/H_sapiens/annotation/GRCh38_latest/refseq_identifiers/GRCh38_latest_protein.faa.gz’). Drug targets and essential genes were aligned with the human proteome using blastp with <=35% identity and 0.0001 e-value. Further, octamer search using *in-house* PERL algorithm was done on sequences with no human homologs. Among the prioritized protein list, the core proteins were further filtered for invariance using Clustalw (35). snp-sites tool (https://github.com/sanger-pathogens/snp-sites) (36) was used to get the VCF file of isolates which was later evaluated for variant and invariant protein sequences that were later segregated and prioritized.

### 6. Gene ontology analysis

OmniBox for Blast2GO software v.2.4.5 available at http://www.blast2go.com/b2ghome. https://www.blast2go.com/ (37) was used. Annotation of all potential drug targets was performed by using default parameters of Blast2GO tool. InterProScan (38) analysis was also done to get the related GO terms. The GO slims and enzyme mapping of potential drug targets using KEGG database (39).

### 7. Web based platform for comprehensive analysis of drug resistant determinants

#### 7.1 Platform architecture

In order to provide an accessible and user-friendly search interface, Ab-AMR is provided as a web-based platform. As can be seen in **Figure 1**, the platform is divided into several sections for ease of navigation. DR determinants can be browsed using their gene name, drug name, mechanism of action, plasmid vs chromosome mediated DR and drug targets can also be browsed using their protein ids, localization and pathways, etc.

**Figure 1.**
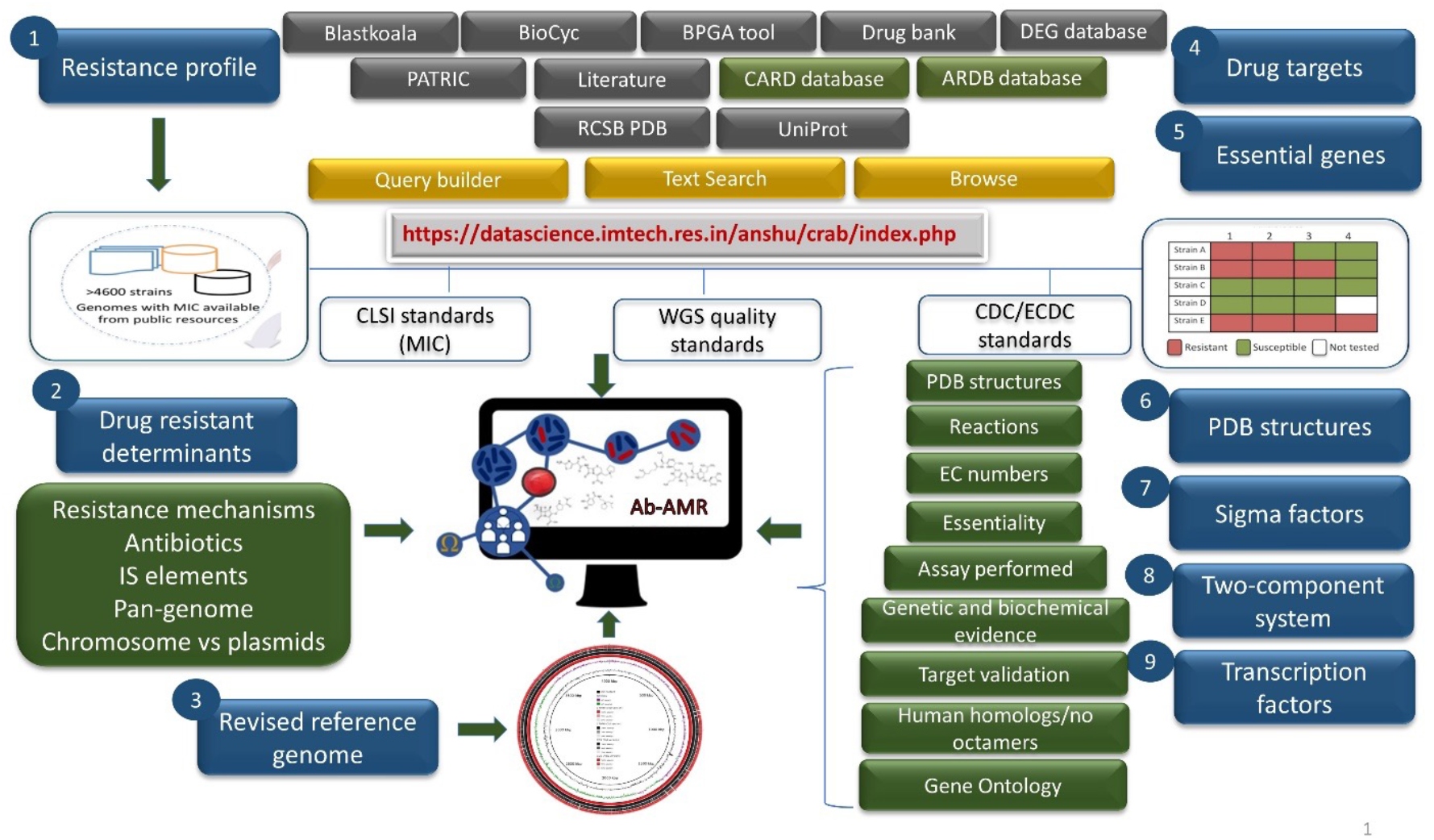
‘Ab-AMR’architecture.

#### 7.2 Utility and search interfaces

Two search options have been provided on the “Ab-AMR” web interface namely ‘Search’ and ‘Advanced search (query builder)’. Simple search option allows text-based search for the users on selected fields of Ab-AMR. Some of the major fields are potential drug targets which can be searched by core, along with their pathway and function, locus tag, gene id, gene product, function, protein id, localization, essentiality (virulence/survival), condition of essentiality (*in vivo*/*in vitro*), drug targets (computational/ experimentally predicted), reported inhibitor, transcription factor, sigma factor, two-component system, pathway, protein ID, reported resistance mechanism, reported β-lactamase family, DR determinants of particular antibiotics, DR determinants (Plasmid vs chromosome mediated), etc. The query builder option facilitates querying Ab-AMR through complex queries with the help of logical operators such as AND & OR. This option allows the user to query the database in a flexible manner and refine the results relevant to user requirements. The browsing interface allows easy way to explore the drug resistant determinants with comprehensive annotations.

Ab-AMR is implemented using the open-source LAMP solution stack on Red Hat Enterprise Linux 5 (IBM SAS x3800 machine) with MySQL (5.0.51b) and Apache (2.2.17) in back-end and frontend web interface is implemented with the PHP (5.5.6). It was designed and tested for web browser Firefox ESR 52.7.2 (64 bit) with 24” Dell Inc with 1920 * 1080 resolution.

## 3. Results

### 3.1 Deep annotation of the reference genome (NZ_CP018664)

The reference AB genome with accession-CP000521.1 and the genes of revised reference genome with accession-NZ_CP018664 are obtained from NCBI and mapped using MEGABlast to get the coordinates of the newer version of the genes within the older version of the genome. Out of 3838 genes in the revised reference genome, 3744 genes show exact alignment with the genes of the older version (accession-CP000521.1). Of the remaining 94 genes in the newer version, 86 genes show multiple hits with different coordinates of the older version of the genome. The remaining 7 genes show partial match between the two genomes. The mapping of gene coordinates was done by developing an *in-house* algorithm. The genome statistics can be seen in **Table 1**.

**Table 1.**
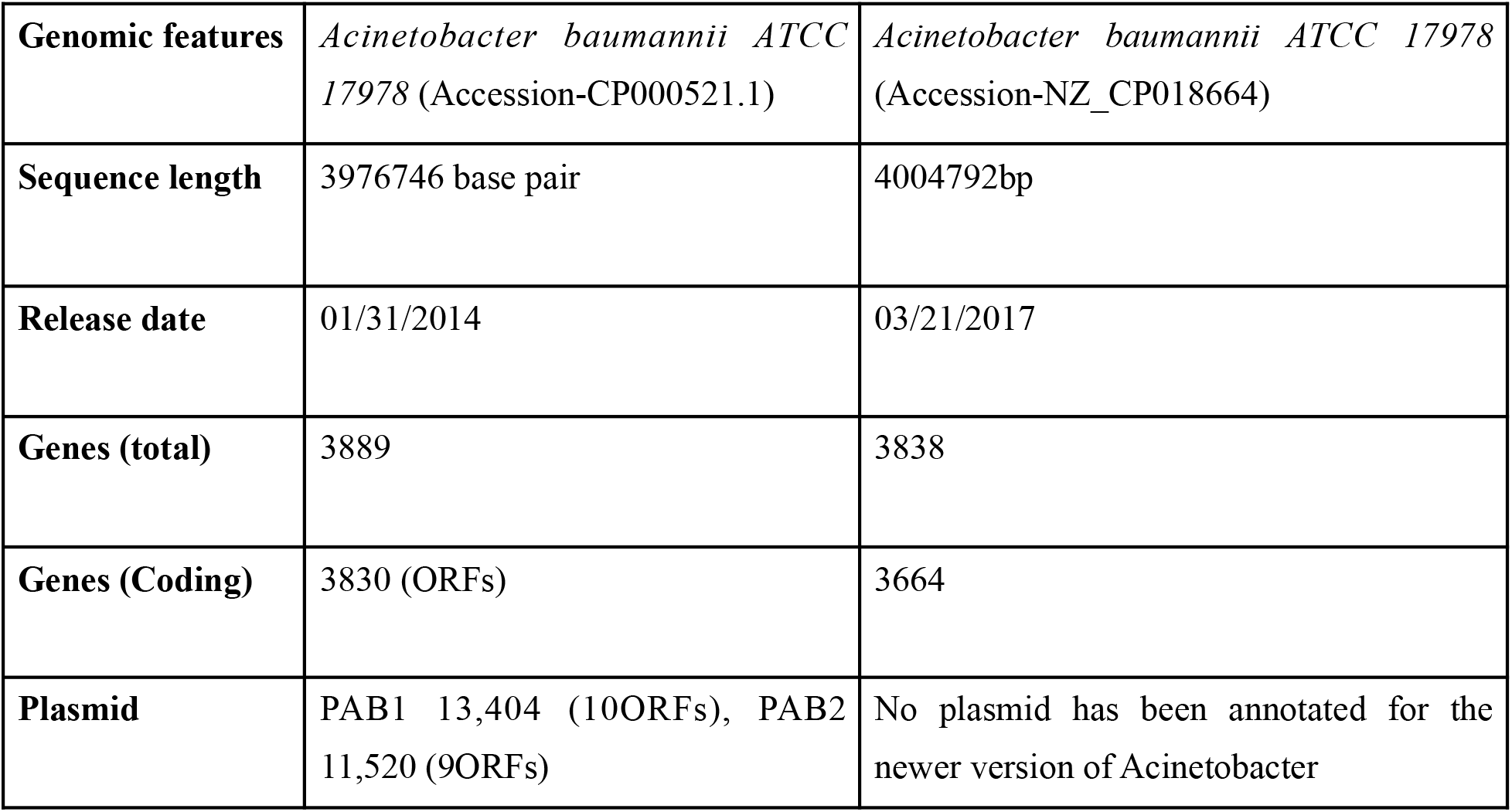
Genome statistics of *Acinetobacter baumannii* ATCC 17978 (accession-CP000521.1) and *Acinetobacter baumannii* ATCC 17978 (accession-NZ_CP018664)

The mapping of gene coordinates was done by developing an *in-house* algorithm. In Ab-AMR, the reference coordinates of the revised reference genome are used for reporting annotations.

The genes were annotated to retrieve gene-protein reactions (GPRs) of AB. For obtaining the pathways data KEGG, BioCyc and PATRIC were searched. Locus tags were mapped with the revised version of reference genome and pathway data was retrieved for 1334 genes. Similarly, data was obtained for the 221 PDB structure using RCSB-PDB. Based on literature scanning, 118 transcription factors, 4 sigma factors and 14 two-component systems are also reported.

A total of 788 genomes were downloaded from NCBI and their proteome was obtained using the RASTtk pipeline. The pan genome of 788 clinical isolates of AB consisted of 2881610 protein sequences, which included 1777728 core (61.69%), 1101485 accessory (38.22%) and 2397 (0.083%) unique genes, respectively. The unique pan genome comprises 12640 protein sequences including the total number of unique core genes as 2256 with an open pan genome (**Figure 2**).

**Figure 2.**
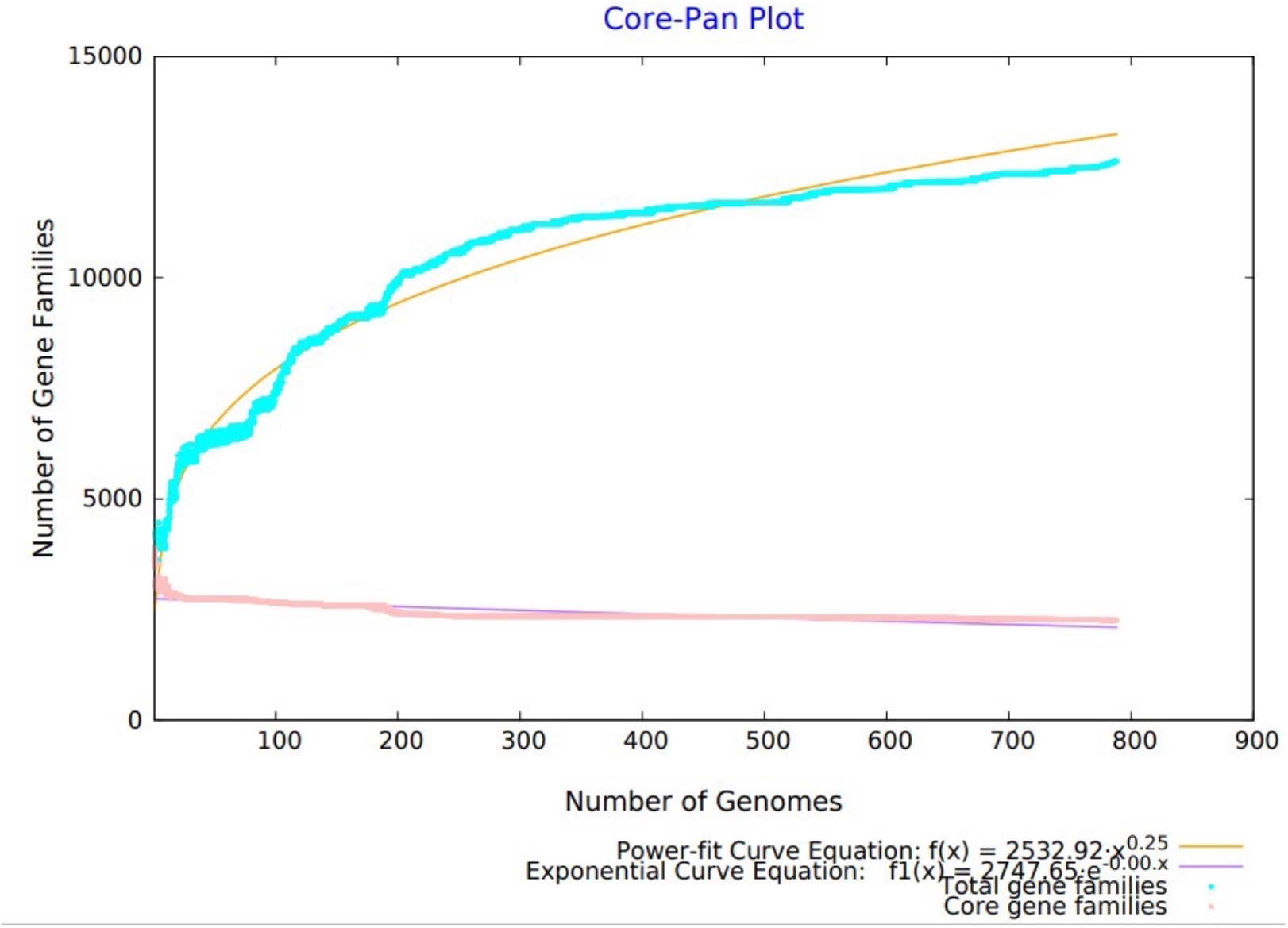
Pan-core plot of 788 AB genomes. The X and Y axis represent the number of genomes and the number of gene families respectively. The pangenome curve which is shown in brown color is not flattened which clearly indicates that AB has an open pangenome.

According to pan-genome calculations, the value of b i.e., 0.248076 in the power-law regression model is the indication of a nearly close pangenome for *AB*.

Based on literature mining, 81 drug targets were curated. The literature was searched for computational and biochemical evidence, whether the target is validated *in vitro or in vivo*, etc., and it was found that 69 out of 81 drug targets were computationally predicted while 12 are experimentally validated (**Figure 3A**). In addition, essential genes were also retrieved from literature and DEG and 614 essential genes were retrieved. The literature also searched for the essentiality of the gene product (survival and virulence), reported inhibitor, condition of essentiality (*in vivo* & *in vitro*), assay/technique performed and criteria of essentiality. Out of 614 essential genes, 424 were essential for the survival of AB and 142 were essential for virulence while the condition of essentiality for 409 showed *in vitro* evidence and 157 showed *in vivo*evidence (**Figure 3A, 3B**).

**Figure 3.**
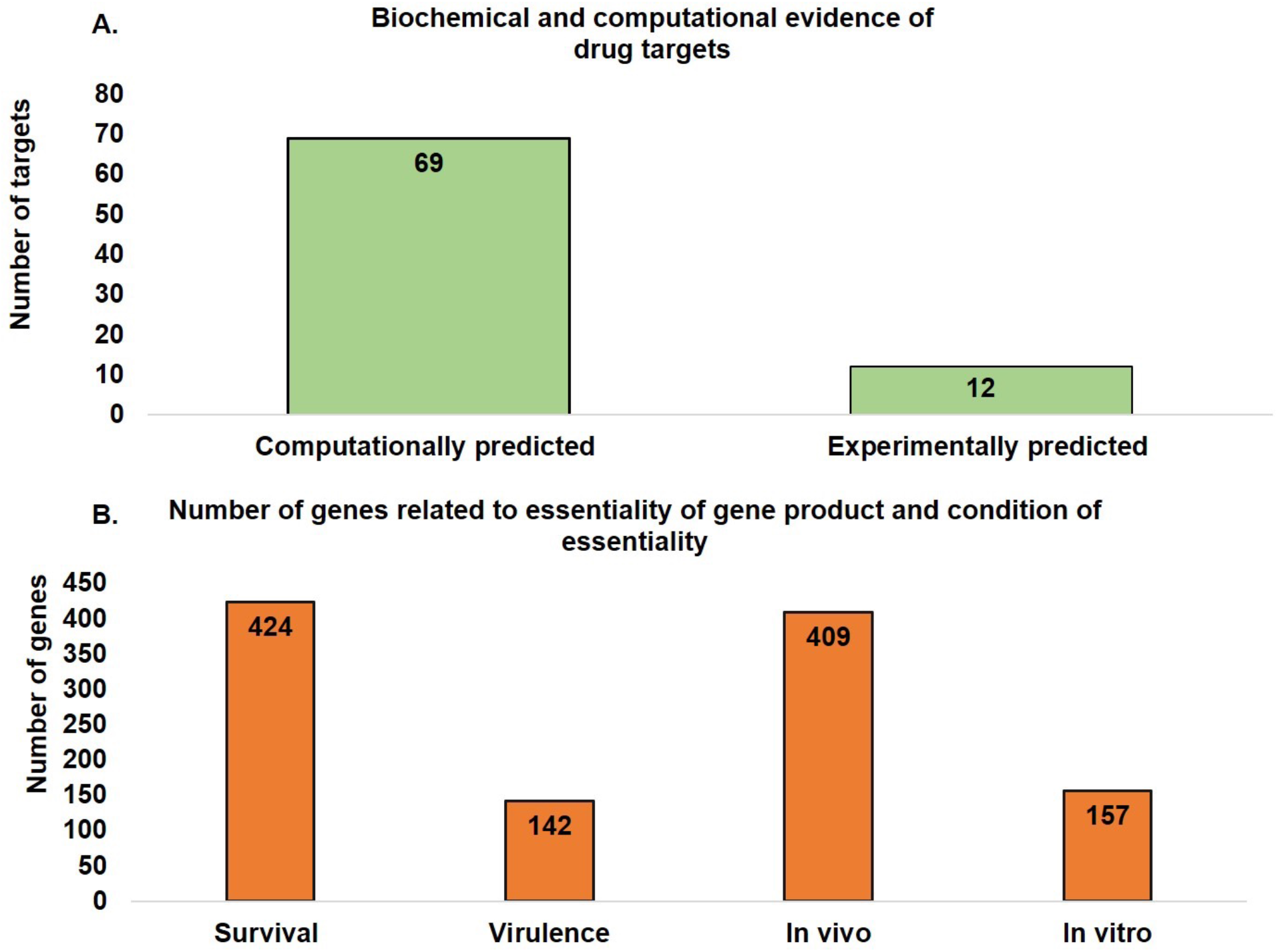
**A**. Genetic and Computational evidence of 81 drug targets. **3B**. Essentiality of gene product (Survival and Virulence and Condition of essentiality i.e. *In vivo* and *In vitro*.

It was observed that 58, out of 81 drug targets are also reported as essential genes. A union of these 58 along with 614 essential genes leads to a total of 637 potential drug targets. These 637 candidates are further analyzed to report the list of most promising drug targets in AB. These 637 proteins are used for subtractive proteomics with human proteome based on two criteria - 1) blastp search with percentage identity <= 35% and e-value 0.0001 and 2) Octamer search (using *in-house* PERL algorithm). It was observed that out of 637, 550 proteins dont show significant alignment with the human proteome. Therefore, total 550 proteins were subjected to octamer search and 261 proteins dont show an octamer match with the human proteins. Out of these 261 proteins, 238 proteins belong to the core proteome of AB and are present in all 788 genomes. The invariant analysis was performed on 238 core proteins and 68 invariant proteins were identified (**Figure 4**).In order to eliminate literature bias in prioritizing drug targets, an unbiased approach is independently applied on the entire proteome of AB using similar steps and 94 invariant proteins are identified as potential drug targets from this strategy too and this prioritized set is then further evaluated for enriched functions and pathways in AB.

**Figure 4.**
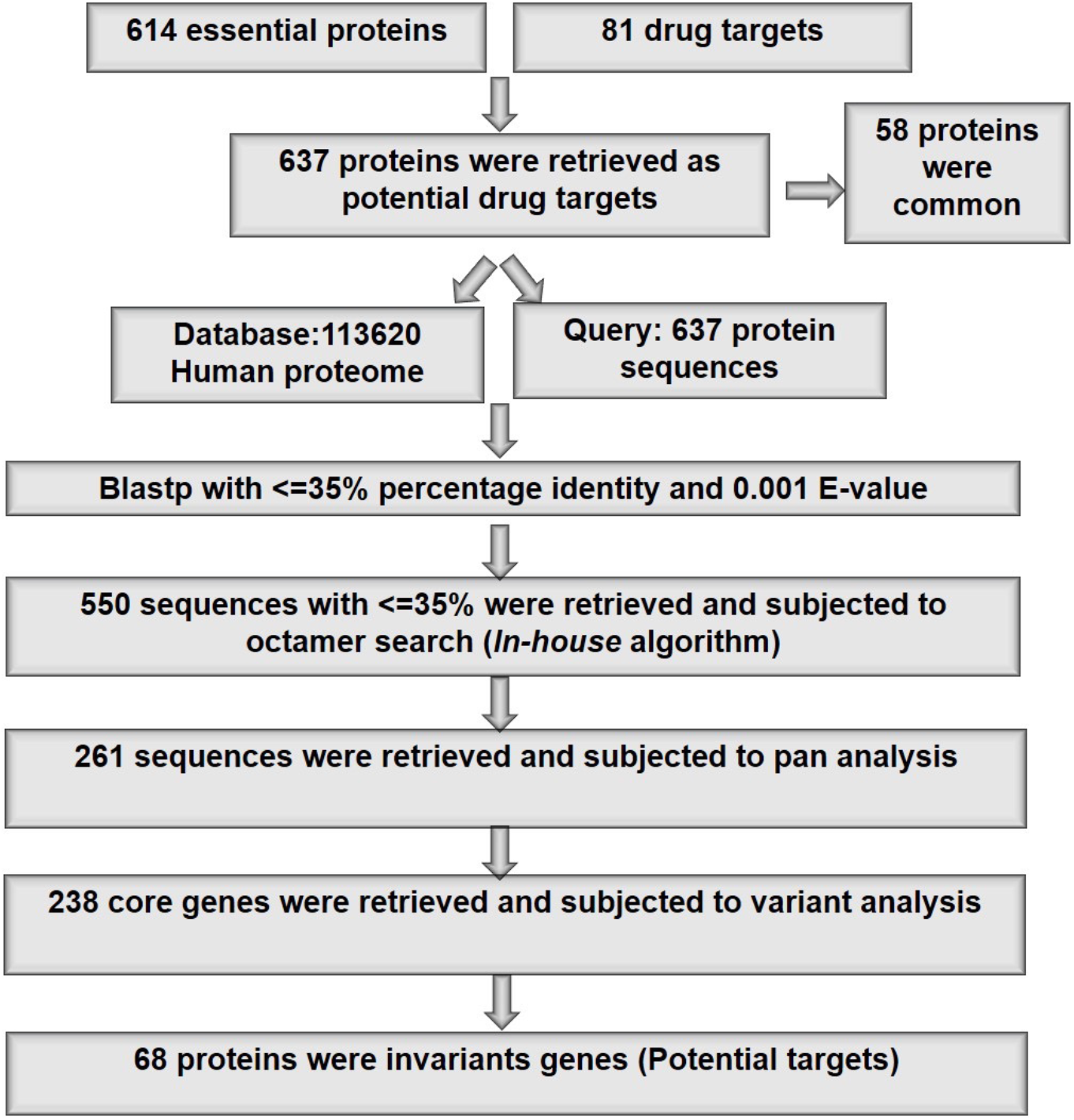
Subtractive proteomics approach to prioritized drug targets

After implementing blast2Go analysis, it was found that the prioritized drug targets are involved in many essential biological functions. Blast2go was used to analyze molecular, biological and cellular function distribution among 68 proteins and it was observed that most of the proteins are the structural constituent of ribosomes (molecular functions). At least 16 proteins belong to translation while other proteins belong to other biological processes which can be seen in **Figure 5 and Figure 6**.

**Figure 5.**
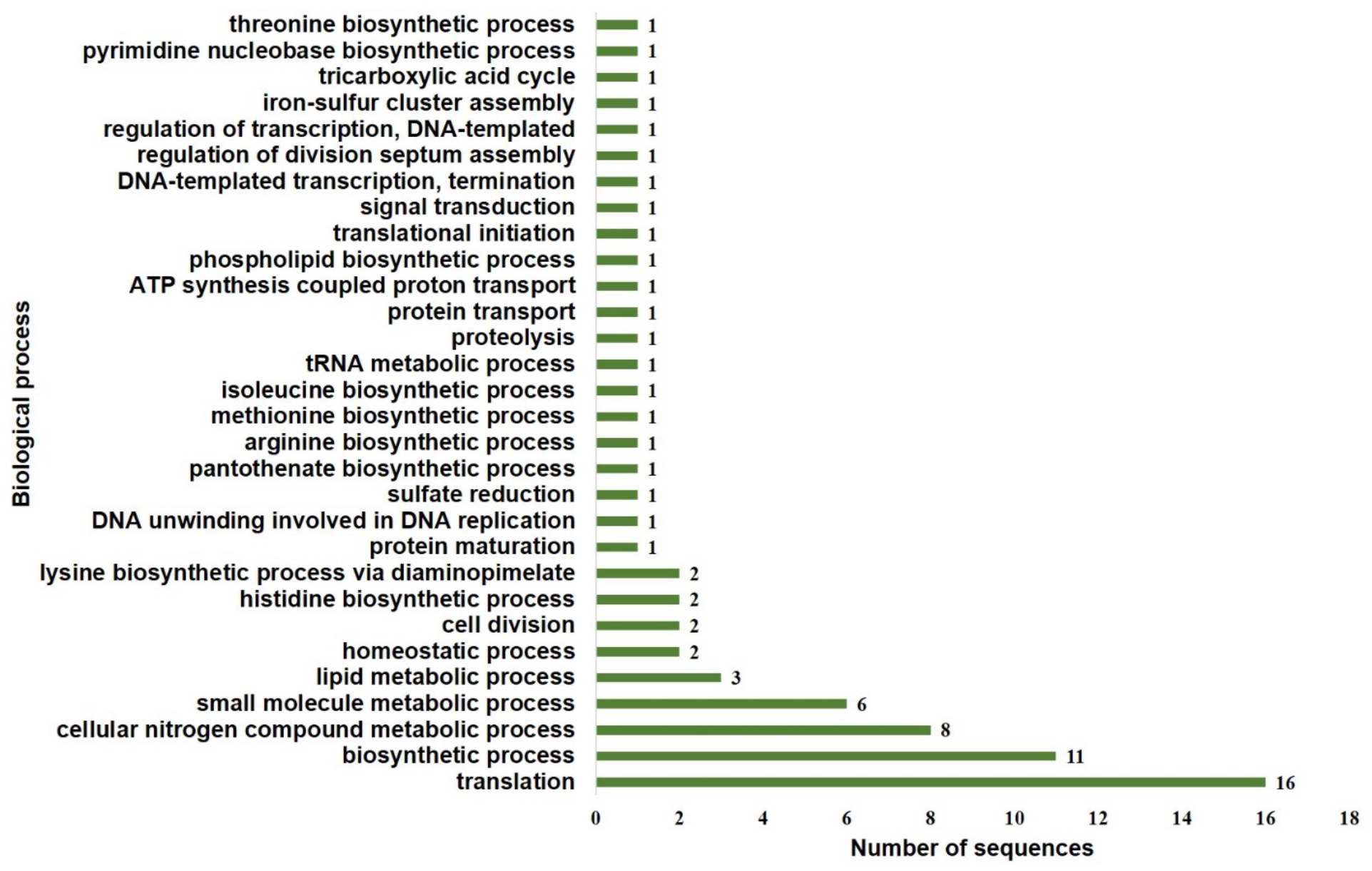
Biological process of 68 therapeutic drug targets.

**Figure 6.**
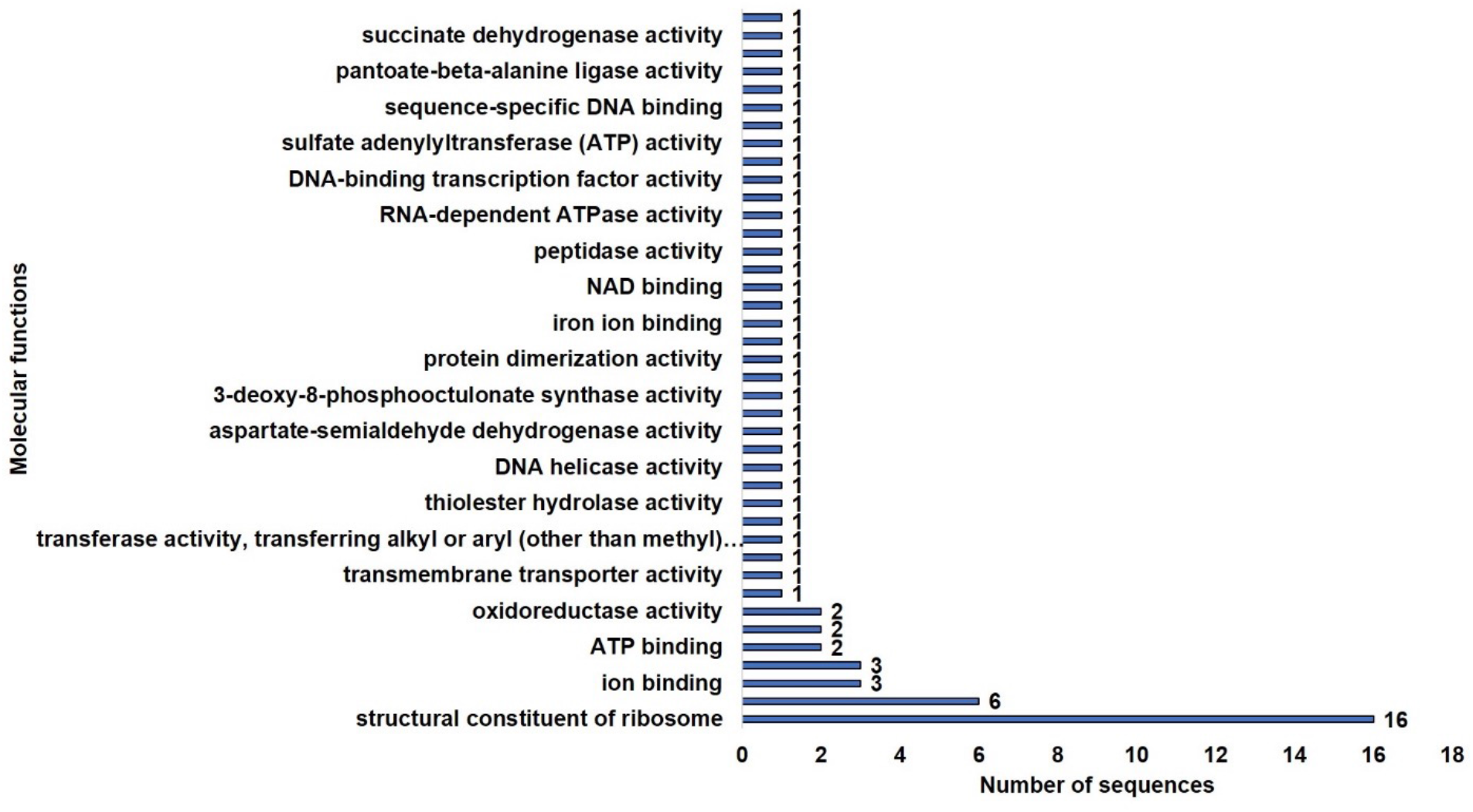
Molecular process of 68 therapeutic drug targets

Out of 94 proteins, 16 proteins are hypothetical. The Blast2GO analysis of such proteins showed no results. It seems like these proteins are not annotated in any database yet. Thus, it is suggested that the functional annotation of such proteins should be explored further.

The spread of MDR AB is of particular concern and to combat resistance, the most crucial step is to assemble the resistant determinants at one place that give rise to the phenotype (38). Based on data gathered in Ab-AMR, it is observed that antibiotic inactivation (β-lactamase, aminoglycoside modification, chloramphenicol acetyltransferase), efflux pumps, alteration of target sites and permeability defects are resistance mechanisms which have been reported for AB. Enlisted below is the statistics of reported resistance mechanisms which have been embedded in Ab-AMR (**Table 2**). It was observed that antibiotic inactivation due to β-lactamases is the most common resistance mechanism in AB (**Table 2**).These β-lactamases were distributed among all four molecular classes i.e. class A, B, C and D based on their amino acid sequences and conserved motifs(39), for instances, CARB(40), CTX(41), GES(42), KPC(43), TEM(44), VEB(45), PER(46), SHV(47), SCO belong to class-A, IMP, NDM(48), VIM(49), SIM and TMB belong to class-B, ADC, CMY which belong to class-C, OXA family which belongs to class-D(50) (**Figure 7**).

**Table 2.**
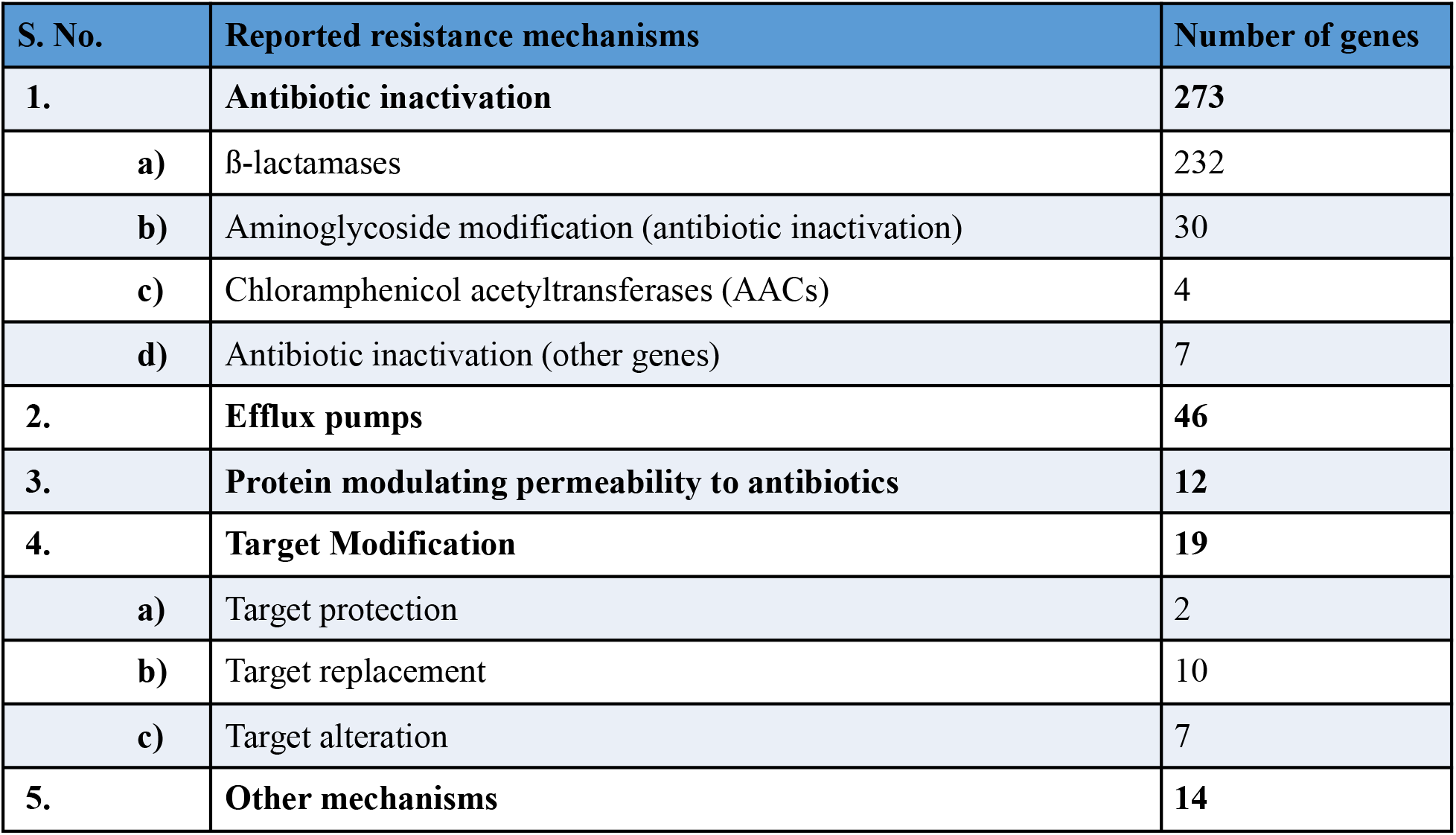
Distribution of resistant determinants in Ab-AMR with reported resistance mechanisms

**Figure 7.**
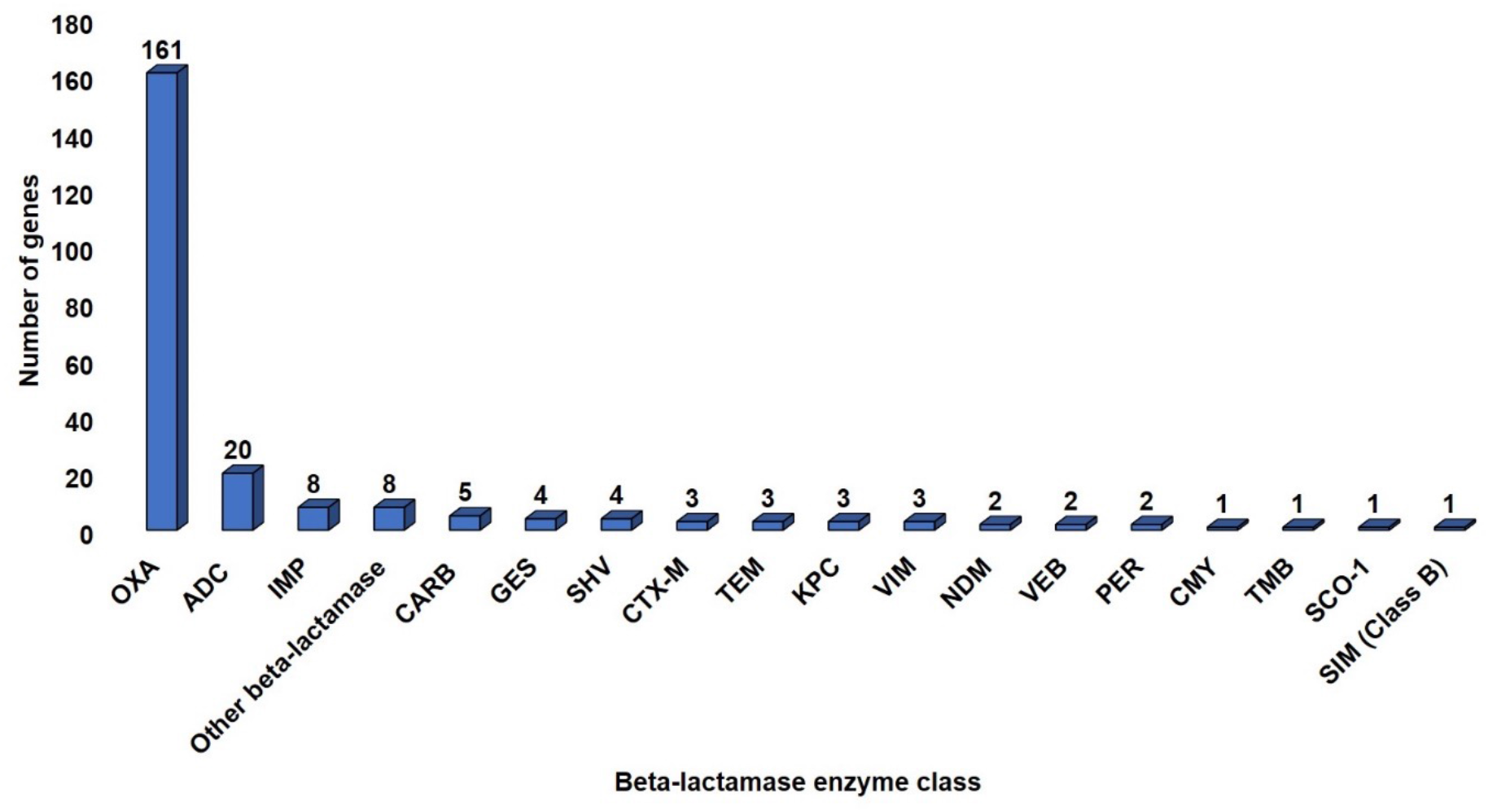
Distribution of beta-lactamase enzyme class among AB genomes. In brackets, beta-lactamase class viz. Class A, B, C and D are mentioned.

The drug resistant determinants were also analyzed whether they are chromosomally mediated or plasmid mediated and surprisingly it was observed that 350 (96%) resistant determinants in AB is associated with the chromosome while only 14 (4%) were plasmid mediated (**Figure 8**).

**Figure 8.**
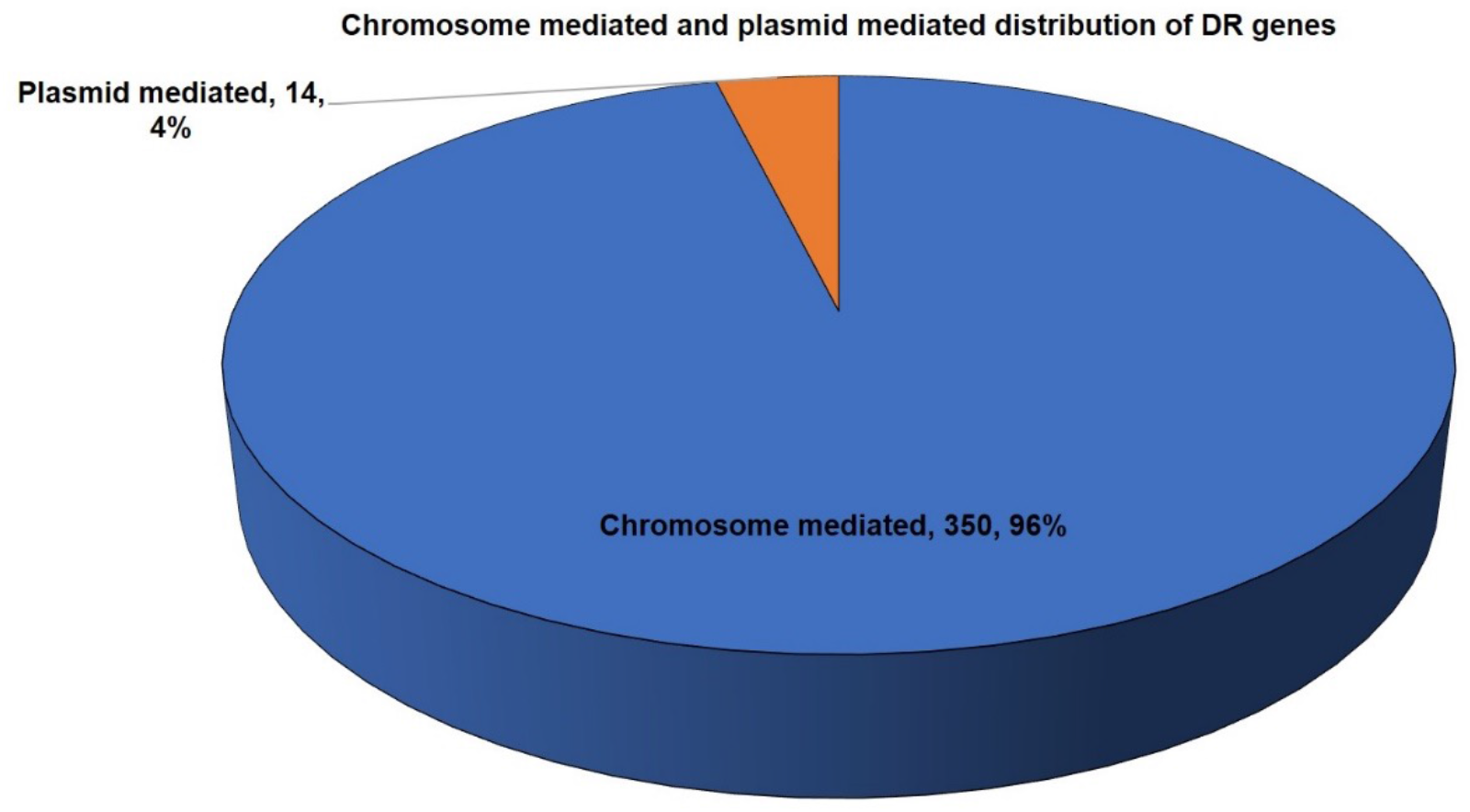
Distribution of plasmid and chromosome mediated DR genes.

The pathway analysis of 364 genes shows that the majority of genes i.e., 31 genes belong to β-lactam resistance while 10 belong to metabolic pathways (**Figure 9**).Out of 364 DR genes, 329 i.e., 90.6% were annotated for their pathways.

**Figure 9.**
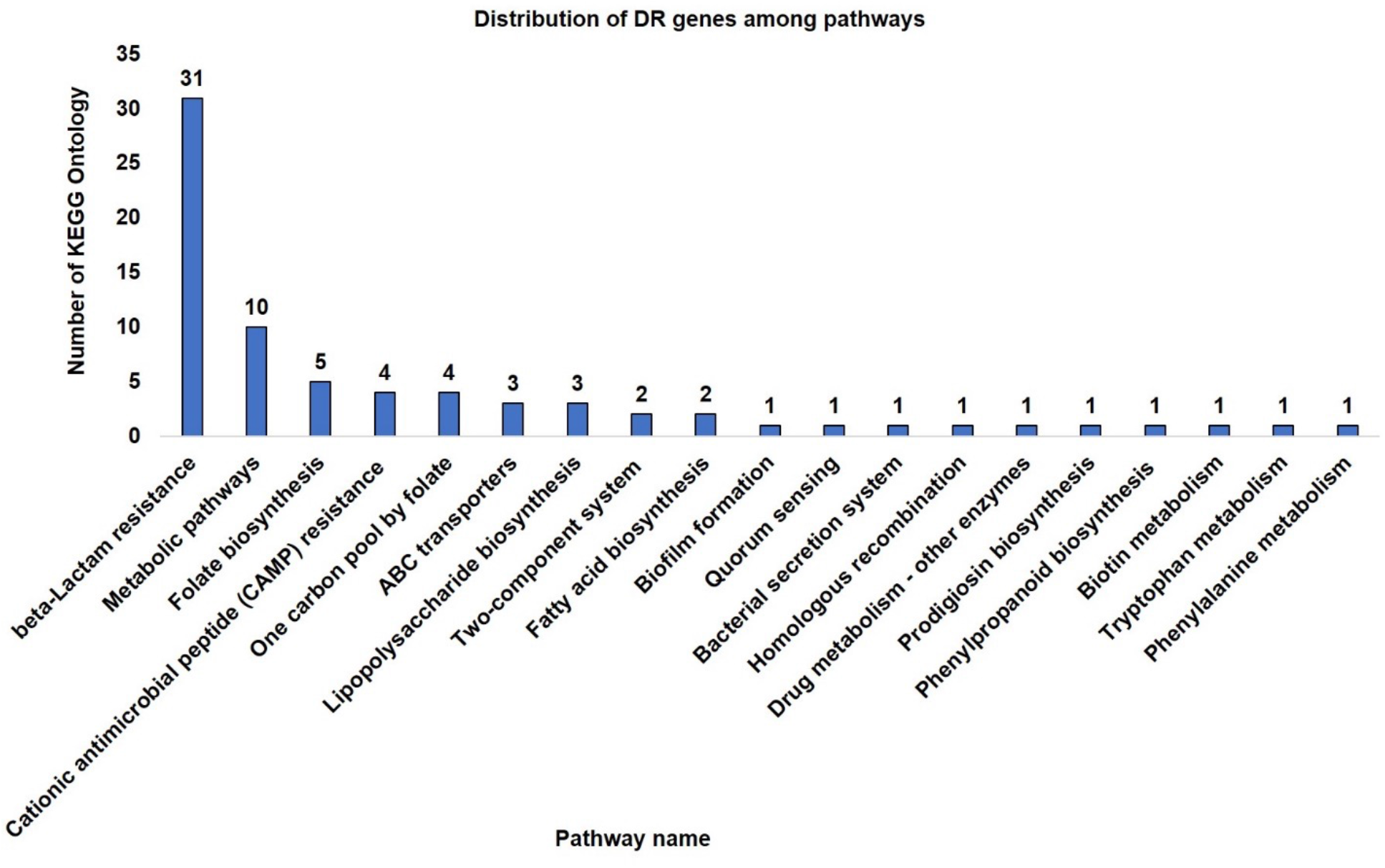
Pathway annotation of 329 drug resistant genes of *Acinetobacter baumannii*.

### 3.2 Mapping of drug resistant determinants to core and accessory genes

Of the 350 drug resistant determinants in the chromosome, 212 (66%) map to the core genome, 111(34%) are accessory while 1 (0%) is unique across 788 genomes of AB (**Figure 10**).

**Figure 10.**
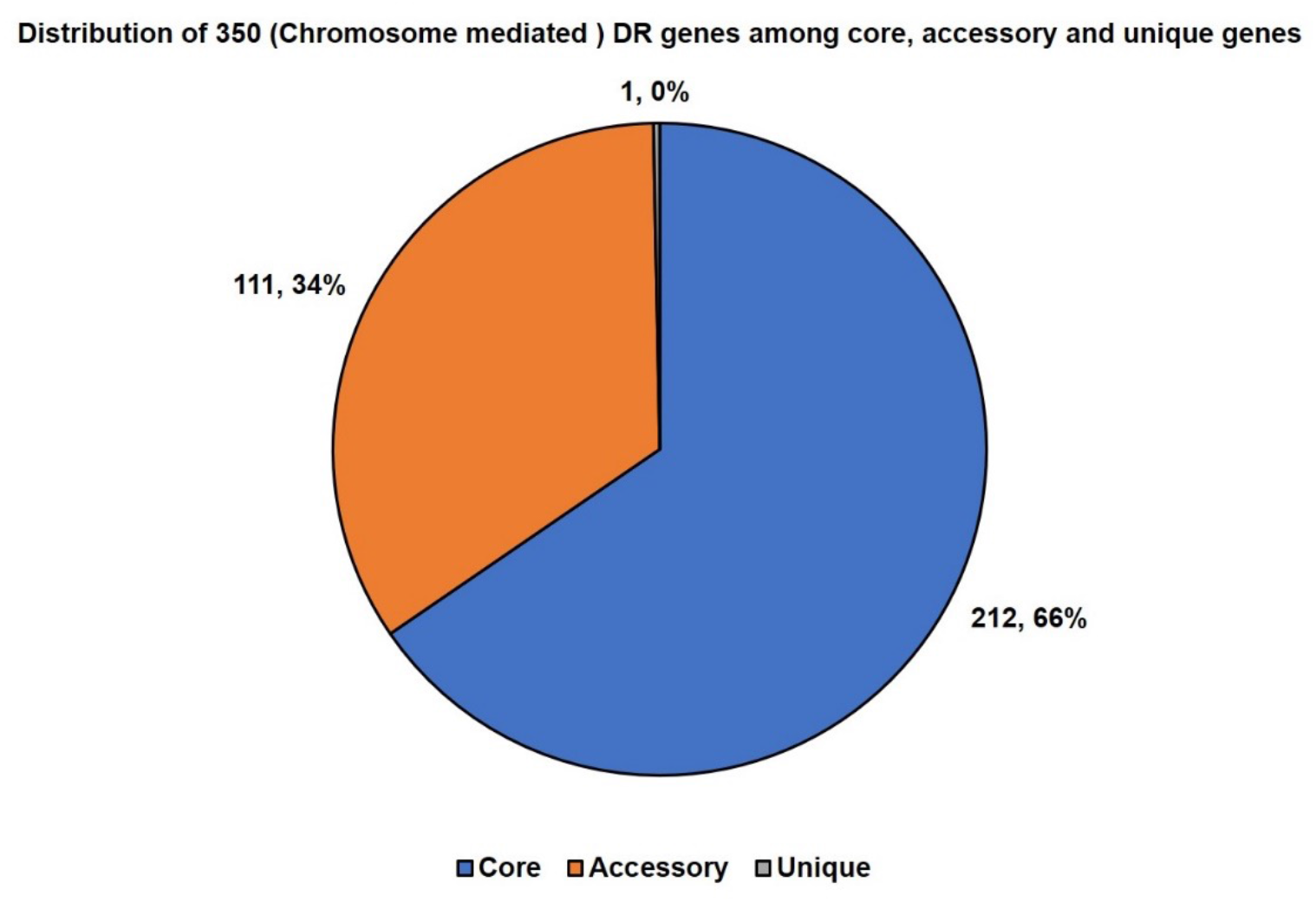
Distribution of 350 chromosome mediated DR determinants among core, accessory and unique.

#### 3.3.5 User-friendly web-based platform: Ab-AMR

The current version of Ab-AMR provides a drug resistance profile of 788 genomes. As of now, 364 DR determinants associated with antibiotic inactivation, efflux, protein modulating permeability and alteration of target site are comprehensively annotated. In addition, data from pan-genome analysis across 788 genomes is also provided for identification of core and accessory DR determinants. Ab-AMR is made using the standard php-mysql framework and offers various search tools including a query builder that facilitates query on over 60 different features for addressing complex questions like core genes which are also essential and have a role to play in drug resistance with no known human homolog, etc.

## 4 Discussion

AMR among AB has increased at a very high rate leading to increased morbidity and mortality and treatment cost in healthcare-associated infections (51). Researchers acknowledged that the terms MDR, XDR and PDR were encountered with various definitions leading to the confusion in the correlation of the existing genotype and phenotype (52). In 2012, A joint initiative by European and United States Centers for Disease Control and Prevention (ECDC and CDC) proposed specific definitions based on the list of antibiotics for characterizing the acquired resistance profile of clinical isolates.For AB, MDR were non-susceptible to >=1 agent in >=3 antimicrobial categories, XDR were non-susceptible to >=1 agent in all but <=2 categories while PDR non-susceptible to all antimicrobial agents listed (54). Prior to this publication, there was no standard definition for the term “MDR”, “XDR” and “PDR for AB (**Table S4**).

The 788 clinical isolates comprising resistance profiles following CLSI standards for MICs and EUCAST standards for whole genome sequence quality and CDC/ECDC standards for acquired resistance profiles were categorized among susceptible, MDR. XDR and TDR isolates using an *in-house* algorithm. It was found that 624 strains were classified as potential MDR. A high number of resistance isolates were observed for amikacin, ciprofloxacin, cefotaxime, ceftriaxone and a combination i.e., trimethoprim-sulfamethoxazole as the number of resistant isolates for these antibiotics was greater than 600. With the existing data of Ab-AMR, multiple resistance genes have been observed for one class of antibiotics for instance target modification and antibiotic inactivation, resistance have been observed for the class of aminoglycosides. Similarly, for macrolide antibiotics, antibiotic inactivation for the mphA gene and antibiotic efflux for msrE genes was observed.

It was observed that the most common resistance mechanism is the antibiotic inactivation (273) genes as most resistant determinants belong to the β-lactamase enzymes (232 genes) followed by aminoglycoside modification enzymes (30), chloramphenicol acetyl-transferase (4) and others antibiotic inactivation enzymes for which exact resistance mechanism is not known. Of 232, 161 β-lactamase genes belong to OXA β-lactamase (class D) enzymes. Based on the observation that out of 364 DR genes, 350 are associated with the chromosome while only 14 genes are on plasmid, it may be indicative of the fact that AB possesses a range of intrinsic DR determinants. Also, of the 350 genes, 212 are in the core genome. The majority of core DR determinants belong to the OXA family which shows resistance via antibiotic inactivation. Other core proteins which belong to protein modulating permeability to antibiotics were associated with the OprD family, OprB family, BenP, CarO, OmpA. For target modifications LpxD, LpxC, LpxA, ParC, GyrA and FabG are reported. For antibiotic efflux mechanism, adeC, abeS, LysR, adeK, aden, abeM, qacE, tolC, emrB, macB, AdeIJK, adeI, macA, craA, and emrE were found. The DR determinants associated with plasmids of AB belong to antibiotic inactivation and antibiotic efflux. These genes are CTX-M-115, StrA, OXA-167, OXA-10, OXA-87, IMP-10, IMP-11, NDM-1, AphA6, aadB, arr-2 which belong to antibiotic inactivation while there are three plasmid mediated genes i.e., msrE, floR and tetA which contribute to antibiotic efflux.

As of now, 364 DR determinants associated with antibiotic inactivation (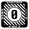-lactamases, aminoglycoside modification, chloramphenicol acetyltransferase), efflux, protein modulating permeability and alteration of target site are comprehensively annotated. In addition, data from pan-genome analysis across 788 genomes is also provided for identification of core and accessory DR determinants. This research work is also focused on the complete genome sequence of *Acinetobacter baumannii ATCC 17978*, a strain which was observed to be significant to study the abundance of genome sequences of *Acinetobacter baumannii* (AB) for their virulence and pathogenesis factors. AB *ATCC 17978* has been used as a laboratory reference strain. Based on synteny maps it was observed that accession CP000521.1 and NZ_CP018664.1 have differences in genome structure and therefore coordinate remapping was performed with NZ_CP018664 version of the reference genome of AB.

Ab-AMR is made using the standard php-mysql framework and offers various search tools including a query builder that facilitates query on over 60 different features for addressing complex questions like core genes which are also essential and have a role to play in drug resistance with no known human homolog, etc. Ab-AMR offers a centralized data resource for systematic mapping of DR determinants, both plasmid and chromosomal mediated, along with deep annotation of clinical isolates. After retrieving ~800 clinical isolates following global standards using Galaxy ASIST and systematic analysis of reference genome of AB ATCC 17978, the Ab-AMR repository was developed in the second objective, in order to ensure that the datasets have relevance both to the research and clinical community. Here, in this research work, we have presented the first version of ‘Ab-AMR’ with structured information of resistance profile of clinical isolates with their available antibiogram i.e. MICs (according to CLSI and classified into their acquired resistance profile as suggested by ECDC and CDC i.e. susceptible, MDR, XDR, TDR, comprehensive annotation of drug resistant determinants as well as genome annotation of revised version of reference genome i.e. *Acinetobacter baumannii ATCC 17978* which will provide rich description of AB in terms of drug targets, essential genes, DR determinants, sigma factors, transcription factor, two-component system etc. Therefore, the first version of ‘Ab-AMR’ has been developed to keep those data fields in mind that will help users in understanding the complexity of the AB genome and its resistance mechanism as it provides the fully structured details of AB clinical isolates. The genotypephenotype analysis of the data using ‘Ab-AMR’ remains to be automated which will hopefully be done in the second version of ‘Ab-AMR’ via which a large number of automations in terms of understanding genotype-phenotype correlation can be made using a single hit.

## 5. Conclusion

The imprudent use of antibiotics has already led to the development of MDR and XDR isolates of AB. Although there exist a number of repositories, software and tools for the prediction of resistome among pathogens, Ab-AMR is the first *Acinetobacter baumannii* repository for understanding the molecular determinants of resistance. This unique AB specific resource is enriched with detailed annotations of drug resistant determinants and houses the largest set of WGS data as per the globally accepted standard for AB till date. The deep annotation of clinical isolates of AB led to the identification of potential new drug targets and a comprehensive landscape of drug resistant determinants.

## 6. Availability of database

This database is freely accessible at https://datascience.imtech.res.in/anshu/ab-amr/index.php (CC BY-NC-SA). The mastersheet, input, output and a program in perl for genome comparison is available at: https://github.com/tinabioinfo/Comparision_of_reference_genome

